# CoREST Complex Stabilizes MYC Protein to Promote Cancer Cell Genome Stability

**DOI:** 10.1101/2024.12.23.630174

**Authors:** Abdul A Khan, Soojin Kim, Tabinda Sidrat, Jaemin Byun, Ariel A Aptekmann, Piero D Dalerba, Johannes L Zakrzewski, JiYeon Shin, Byungwoo Ryu

## Abstract

The CoREST complex is a multi-subunit epigenetic regulator implicated in histone modification and transcriptional repression, but its role in tumorigenesis is not well-defined. Here, we show that the CoREST complex directly interacts with and stabilizes the MYC oncoprotein in cancer cells through site-specific deacetylation of lysine residues, primarily mediated by HDAC1/2.

These modifications protect MYC from proteasomal degradation independently of transcriptional regulation, maintaining high MYC protein levels in cancer cells. Transcriptomic analysis reveals that the CoREST-mediated MYC stabilization activates transcription of genes critical for DNA replication and mitotic chromosome segregation, and enhances melanoma cell viability. These findings suggest that the CoREST complex maintains cancer cell genome stability and promotes survival by sustaining MYC oncogenic activity, highlighting it as a potential therapeutic target in MYC-driven malignancies.

## Introduction

The CoREST (Corepressor of REST) complex is a multi-subunit protein complex known for its role in epigenetic regulation, primarily through histone modification and transcriptional repression (*1*). This complex is composed of three core proteins, including histone deacetylase 1 or 2 (HDAC1/2), lysine-specific histone demethylase 1A (LSD1/KDM1A), and REST corepressor 1 (RCOR1) which serves as a scaffold protein connecting the two histone-modifying enzymes (*2, 3*). While initially identified for its function in neuronal development, the CoREST complex has been implicated in a variety of cellular processes beyond the nervous system, including cancer (*4–6*). Notably, CoREST possesses a unique combination of enzymatic activities, containing both histone demethylase (LSD1) and histone deacetylase (HDAC1/2) enzymes, which allows it to modulate chromatin structure through the removal of both methyl and acetyl groups from histone tails(*7*). While traditionally recognized for its role in repressing neuronal genes in non-neuronal tissues through histone modifications, emerging evidence suggests that the CoREST complex may have broader functions in cancer development and progression (*5, 8*).

Histone post-translational modifications (PTMs), such as acetylation and methylation, also play a crucial role in genome stability in cancer cells. These PTMs affect chromatin structure, influencing the recruitment of DNA repair proteins, cell cycle regulation, and ultimately preventing genome instability (*9*). For instance, LSD1/KDM1A partners with RNF168 at DNA damage sites to remove methyl groups from H3K4me2, while genotoxic stress induces deacetylation at H3K9 and H2K56 (*10, 11*). Treatment of HDAC inhibitors also disrupts DNA damage repair in cancer cells (*12*). These experimental results suggest that specific histone deacetylation and demethylation are required for restoring chromatin post-damage. Non-histone PTMs also influence cancer cell genome stability by impacting various cellular processes, including DNA damage response (DDR) by acetylation of ATM activating ATM kinase activity (*13, 14*). Furthermore, a recent report shows that the CoREST complex is recruited to sites of DNA double-strand breaks (DSBs) via GSE1-bound deubiquitinase USP22 (*15*). This finding highlights the critical role of the CoREST complex in facilitating DNA damage repair by creating a favorable chromatin environment that is conductive to DNA repair signaling. As a result, the CoREST complex helps protect the cancer cell genome.

Dysregulation of MYC family oncogenes is a hallmark of more than 50% of human cancers and is frequently associated with poor prognosis and aggressive disease progression (*16*). As a master transcription factor, MYC orchestrates a vast network of genes involved in cell growth, proliferation, metabolism, and apoptosis (*17, 18*). The stability of the MYC protein is tightly controlled through the ubiquitin-proteasome system, and its levels are modulated by various PTMs (*19, 20*). While post-translational phosphorylation is a well-documented major regulatory mechanism of MYC stability(*21*), acetylation of specific lysine residues has also been shown to play crucial roles in determining MYC protein stability and transcriptional activity (*22, 23*).

However, the precise mechanisms and enzymes responsible for regulating MYC acetylation status in cancer contexts remain incompletely understood.

MYC also plays a complex role in the DNA damage response (DDR) and the maintenance of genome stability in cancer cells. Although oncogenic MYC often induces DNA damage through the increased replication stress and reactive oxygen species (*24, 25*), it has also been suggested that MYC transcriptionally activates genes encoding key intermediates of DDR and chromosome maintenance, helping cancer cells survive and maintain genomic integrity under oncogenic stress (*26*). While both the CoREST complex and MYC have been independently studied in cancer, the potential functional connection between these two important regulators has not been previously explored. In this study, we present evidence that the CoREST complex directly interacts with and stabilizes the MYC protein through site-specific deacetylation. As a result, elevated expression of oncogenic MYC leads to the transcriptional activation of genes involved in DDR and mitotic chromosome segregation in cancer cells.

Our study provides new insights into how the epigenetic regulator CoREST complex can influence cancer cell behavior beyond its canonical roles in histone modification and transcriptional repression. Furthermore, it identifies the CoREST complex as a potential therapeutic target, particularly in cancers driven by MYC overexpression, offering new avenues for cancer treatment strategies.

## Results

### Stabilization of MYC protein by physical interaction with CoREST complex in cancer cells

Given the recruitment of the CoREST complex to DSB sites for post-damage histone modifications and MYC’s role in maintaining cancer cell genome stability, particularly through transcriptional activation of genes mediating DDR, we investigated whether the CoREST complex regulates MYC expression in cancer cells. First, proximity ligation assays (PLAs) revealed that MYC and each core component of the CoREST complex (HDAC1/2, RCOR1, LSD1) are in close proximity exclusively within cell nuclei (Fig. 1A), supporting the likelihood of direct interaction. Co-immunoprecipitation (co-IP) assays further confirmed this interaction *in vitro* and in cells. Recombinant His-tagged full-length MYC was incubated with lysates from MeWo melanoma cells, and reciprocal co-IPs using anti-MYC and anti-RCOR1 antibodies demonstrated robust binding (Fig. 1B). Consistent with this, co-IP assays in HEK293T cells expressing FLAG-tagged MYC and HA-tagged RCOR1 also validated the interaction (Fig. 1B). Furthermore, LSD1 MYC co-IPed MYC in both *in vitro* and in cellular experiments, confirming that MYC interacts with the entire core CoREST complex.

**Fig. 1.**
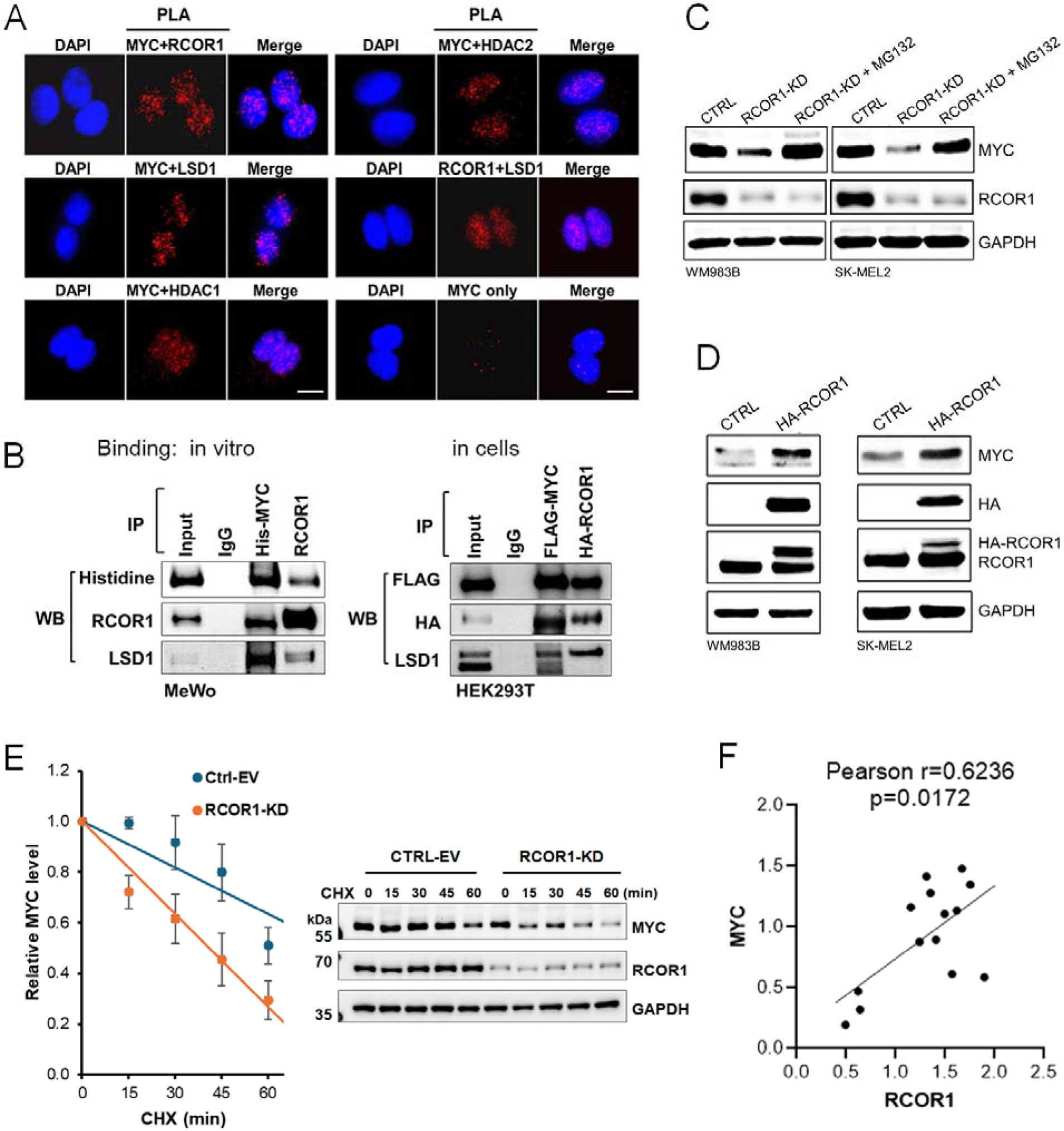
MYC binding to CoREST-complex stabilizes MYC from proteolytic degradation. **(A)** Proximity ligation assay between MYC and each core component of CoREST-complex (RCOR1, HDAC1/2, and LSD1) in WM983B melanoma cells. **(B)** Co-immunoprecipitation assays showing MYC interaction with CoREST complex. *In vitro* binding was performed by incubating purified His-tagged full-length MYC protein with MeWo cell lysate. MYC binding in cells (HEK293T) was performed by expressing FLAG-tagged full-length MYC and HA-tagged RCOR1. **(C)** Immunoblots of MYC levels in melanoma cells with RCOR1-KD and proteasome inhibitor MG132 treatment (10 µM, 4 hrs). **(D)** Immunoblots of MYC in cells exogenously expressing HA-tagged RCOR1. **(E)** CHX pulse-chasing on MYC levels in melanoma cells (WM983B) with RCOR1-KD. **(F)** Positive correlation in expression levels between RCOR1 and MYC proteins in a panel of 14 melanoma cell lines.

Although it remains unclear which core component of the CoREST complex directly interacts with MYC, our mapping studies using FLAG-tagged MYC domain deletion mutants suggested that the DNA-binding C-terminal region harboring a basic-helix-loop-helix leucine zipper (b- HLH-LZ) domain is essential for interaction with the CoREST complex, as all mutants lacking the C-terminal region showed no or limited binding affinity (fig. S1). Previous studies by others have shown that MYC binding domain III (MBIII) interacts with histone-modifying complexes like HDAC3 and SIN3 (*27, 28*); however, the N- and C-terminal truncated central region containing only the MBIIIb and MBIV domains showed no binding affinity with RCOR1.

Conversely, full-length MYC with both MBIIIb and MBIV domain deletions, or with individual deletions of either domain, retained binding affinity comparable to wild-type MYC (fig. S1B). Collectively, these deletion analyses strongly suggest that the C-terminal domain of MYC is critical for CoREST complex interaction.

We also found that the CoREST complex stabilizes MYC protein, protecting it from proteolytic degradation in cancer cells. Knockdown (KD) of RCOR1 reduced MYC protein levels. This effect was reversed by treatment with the proteasome inhibitor MG132 (Fig. 1C). Furthermore, exogenous expression of HA-tagged RCOR1 (HA-RCOR1) in WM983B and SK-MEL2 melanoma cells significantly increased MYC protein levels (Fig. 1D). Cycloheximide pulse-chase analysis of MYC protein in A375 melanoma cells with RCOR1-KD revealed an increased degradation rate compared to isogenic control cells (Fig. 1E), suggesting that the CoREST complex is involved in the dynamic regulation of MYC protein stability in cancer cells. This notion is further supported by positive correlations between MYC and RCOR1 protein expressions observed across a panel of fourteen melanoma cell lines (Fig. 1F).

These findings indicate that PTMs of MYC mediated by HDAC1/2 or LSD1 within the CoREST complex plays a critical role in regulating MYC homeostasis. This PTM-mediated MYC stability is also supported by our findings that RCOR1-KD did not significantly alter *MYC* transcript levels (Fig. 2A), while expression of known MYC target genes (*CCNA2*, *MAD2L1*, *MCM4*, *RRM1*, and *TYMS*) was significantly reduced (Fig. 2B). We also found that melanoma cell growth rate is significantly dependent on the levels of RCOR1. Disruption of CoREST complex-MYC axis by RCOR1-KD in A375 cells resulted in significantly decreased cell viability compared to the isogenic control cells (Fig. 2C). Genetic KD of RCOR1 or LSD1 and pharmacologic inhibition of the CoREST complex by Corin, a dual-action inhibitor of HDAC1/2 and LSD1 (*2*), induced an increased apoptotic cell fraction, an effect also seen with MYC depletion (Fig. 2D). These findings suggest that the CoREST complex promotes oncogenesis, at least in part, by increasing MYC protein levels.

**Fig. 2.**
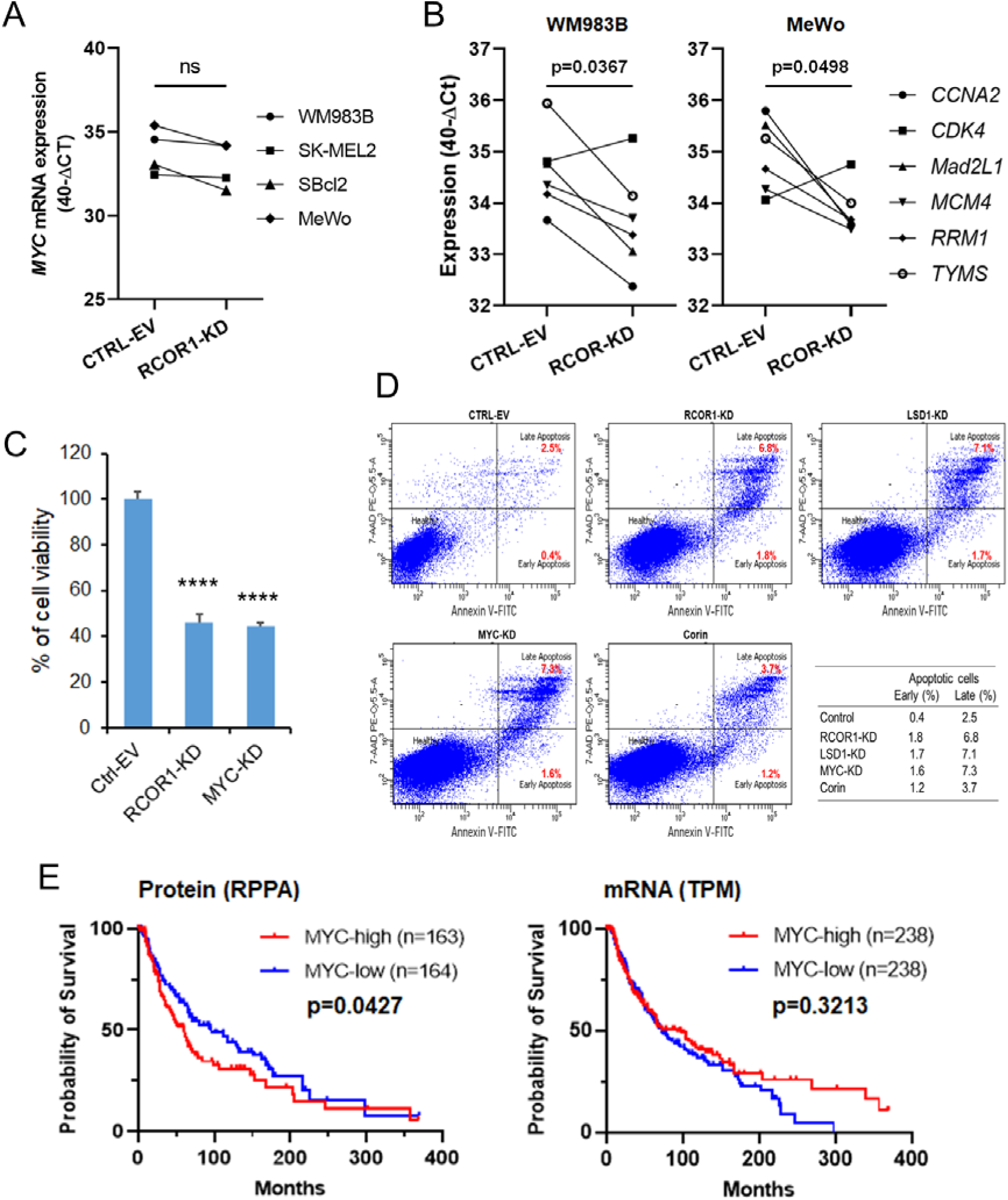
The CoREST complex promotes melanoma cell growth via increased MYC expression. **(A)** RT-PCR analysis of MYC transcript levels in four melanoma cell lines with RCOR1-KD compared to those of control cells. **(B)** RT-PCR analysis of six MYC target genes selected from Hallmark MYC Targets V1 gene set (MSigDB) in RCOR1-KD and control cells. **(C)** Viability assay of A375 melanoma cells with RCOR1-KD and MYC-KD. Mean with s.d., Student’s t-test, 2-sides, ****p<0.0001. **(D)** Apoptosis assays using flow cytometry of WM983B cells with genetic knockdown and pharmacologic inhibition of the CoREST complex. **(E)** Kaplan-Meier survival curves of melanoma patient groups with high and low expressions of MYC protein and transcript levels defined by median expression across patients (SKCM TCGA dataset). RPPA (reverse phase protein array) and RNA-seq datasets were used for MYC protein and mRNA expressions, respectively.

To assess the pathological significance of increased MYC protein stability in melanoma, we analyzed the overall survival of melanoma patients stratified by high versus low MYC expression, at either the protein or mRNA level, using Kaplan-Meier survival curves. Patient stratification based on MYC protein levels showed a significant inverse association with survival, while no association was found with MYC mRNA levels. This finding suggests that CoREST-mediated MYC protein stabilization might play an important role in contributing to the poor survival outcomes of melanoma patients (Fig. 2E).

### HDAC2 in the CoREST complex regulates MYC stability via site-specific lysine deacetylation

To determine which enzyme in the CoREST complex (HDAC1/2 or LSD1) plays a role in the PTM-mediated MYC stabilization, we treated WM983B cells with MS275 (Entinostat), a class I HDAC inhibitor (HDACi); LSD1 inhibitor (LSD1i), GSK2879552; a combination of MS275 and GSK2879552 (HDACi + LSD1i); and Corin (*2*). We found that Corin was the most effective in decreasing MYC protein levels, while the histone demethylase inhibitor (LSD1i) showed no effect (Fig. 3A). Genetic KD of HDAC2 resulted in decreased MYC protein levels in WM983B cells, while no changes were observed following HDAC1 depletion (Fig. 3B), suggesting that HDAC2 in the CoREST complex specifically contributes to the proteolytic regulation of MYC. Pharmacological inhibition of the CoREST complex by Corin resulted in decreased MYC levels across a panel of 10 melanoma cell lines, further supporting MYC stabilization by the CoREST complex via protection from PTM-mediated proteolytic degradation (Fig. 3C). Furthermore, decreased MYC levels following CoREST complex inhibition (Fig. 3A) closely paralleled the growth inhibition patterns observed in melanoma cell lines (SK-MEL2 and 1205Lu) (Fig. 3D).

**Fig. 3.**
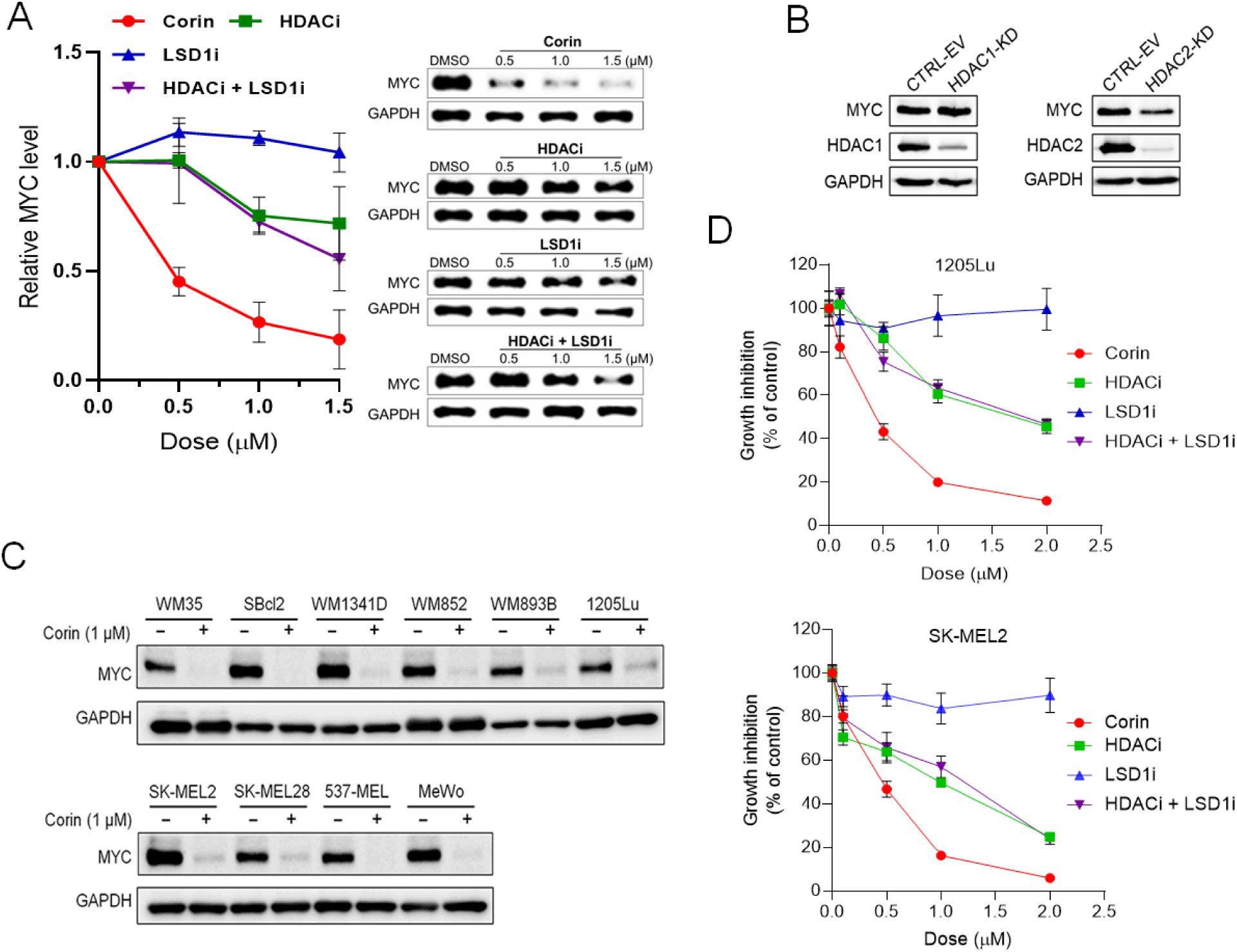
Pharmacologic inhibition of CoREST complex by Corin induces decreased MYC protein expression and melanoma cell growth inhibition. **(A)** Inhibition of CoREST complex by Corin, a dual inhibitor for HDAC1/2 and LSD1, MS275 (HDACi), GSK2879552 (LSD1i), and a combination of MS275 and GSK2879552 (HDACi + LSD1i) in WM983B cells, showing a dose-dependent MYC level decrease by Corin. Cells were treated with the indicated doses of each inhibitor for 36 hrs. Data are the means and s.d. of three independent experiments (right panel). Representative images of MYC immunoblot experiments (left panel). **(B)** Immunoblot analyses of MYC levels in melanoma cells with HDAC1-and HDAC2-KD by shRNA. **(C)** Immunoblot analysis of MYC protein level in melanoma cell lines treated with Corin. All cells were treated with 1.0 µM Corin for 36 hrs. **(D)** Dose-dependent growth inhibition assays with CoREST-complex inhibitor treatments in melanoma cell lines (SK-MEL2 and 1205Lu). Growth inhibition was measured at 72 hrs after indicated doses of each inhibitor treatment.

To identify potential lysine residues targeted by the CoREST complex for deacetylation, we performed site-directed mutagenesis by exogenously expressing mutant forms of MYC with specific lysine-to-arginine substitutions. First, we identified 11 predicted post-translational lysine acetylation residues in MYC using the online tool GPS-PAIL (*29*) and each HA-tagged-mutant form of MYC was expressed in WM983B cells. Next, we treated the cells with Corin and quantified mutant MYC levels by immunoblotting followed by band quantifications using ImageJ analysis (*30*). The quantification of each mutant MYC protein level suggested five lysine residues as potential deacetylation target sites (K148, K157, K275, K289, and K323) of the CoREST complex (Fig. 4A). We then exogenously expressed a mutant form of MYC (MYC-DR) in which all the five lysine residues were substituted by arginine in melanoma cells. The protein level of MYC-DR was higher than that of wild-type MYC (MYC-WT) in both RCOR1-KD and control cells. Importantly, MYC-DR was resistant to proteolytic degradation in RCOR1-KD cells compared to control cells (Fig. 4B). A similar trend was also observed in cells (WM983B and A375) where the enzymatic activity of HDAC1/2 in the CoREST complex was inhibited by Corin treatment (Fig. 4C).

**Fig. 4.**
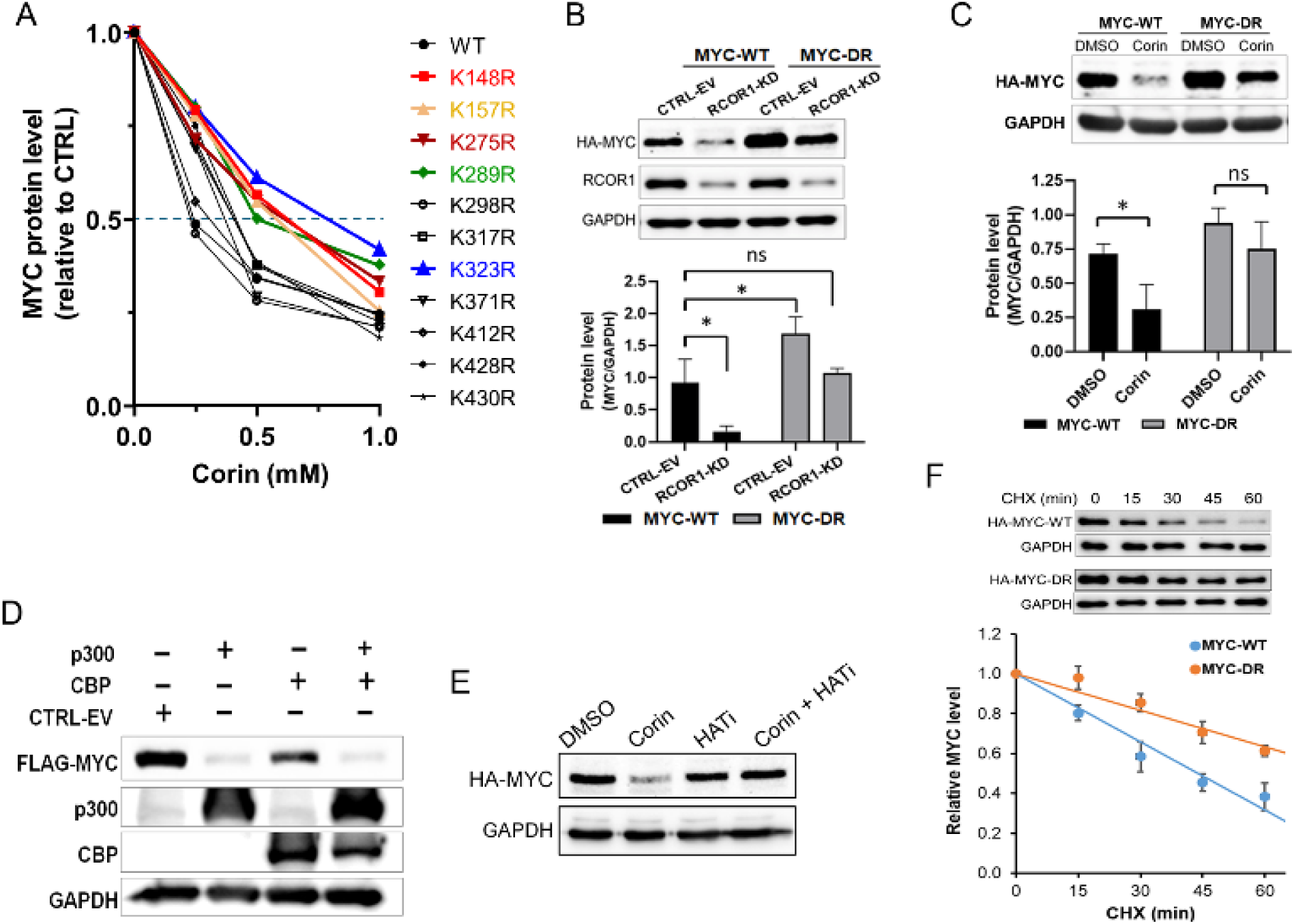
Site-specific deacetylation of MYC by CoREST-complex stabilizes MYC protein in melanoma. **(A)** Site-directed mutagenesis analysis of mutant MYC in WM983B melanoma cells. Each N-terminal HA-tagged mutant MYC at the predicted Lys sites was exogenously expressed in cells and treated with Corin at doses of 0, 0.25, 0.5, and 1.0 μM for 36 hrs. The mutant MYC proteins were immunoblotted with an anti-HA antibody. Quantification analysis of the exogenously expressed mutant MYC proteins remain over 50% at the dose of 0.5 uM of Corin identified as potential Lys residues targeted by HDAC1/2 within the CoREST-complex. **(B)** Mutagenesis assay showing the 5x-mutant MYC (MYC-DR) and wildtype-MYC (MYC-WT) levels in RCOR1-KD and isogenic control cells (WM983B). A representative immunoblots assay (upper panel) and quantification (lower panel) from biological triplicates. Mean with s.d., Student’s t-test, two-sided, * p<0.05. **(C)** Quantification of MYC-DR and MYC-WT measured in a melanoma cell line (A375) treated with 1.0 µM Corin and representative immunoblots from three biological replicates. **(D)** Immunoblots of FLAG-tagged MYC in lenti-X-293T cells with exogenous overexpression of p300 and CBP histone acetyltransferases in relation to empty vector control cells. **(E)** Immunoblots of HA-tagged MYC in WM983B cells treated with vehicle-control (DMSO), Corin (1.0 µM), HATi A485 (3.0 µM), and Corin + HATi combination for 24 hrs. **(F)** Quantification of mutant MYC-DR stability in WM983B cells. HA-tagged MYC-DR and MYC-WT were stably expressed using lentivirus. CHX (10 μM) was added to block cellular protein synthesis at different time points. Representative immunoblots of triplicates (top) and HA-MYC quantification (bottom). The HA-MYC level at each time point is represented relative to the level at time zero.

These data strongly indicate that deacetylation of MYC at these specific lysine residues by the CoREST complex stabilizes MYC and protects it from proteolytic degradation in melanoma by counterbalancing lysine acetylation mediated by histone acetyl transferases (HATs). This notion was supported by the significantly decreased MYC levels in lenti-X-293T cells stably overexpressing p300 and CBP (Fig. 4D). The p300 protein has been shown to regulate MYC expression both transcriptionally and at the post-translational level (*31*). To distinguish from transcriptional regulation of MYC gene expression by p300/CBP, we used WM983B cells stably expressing HA-tagged MYC and treated them with A485, a selective p300/CBP HAT inhibitor (HATi) (*32*). The A485 treatment recovered the MYC protein level that was reduced by Corin treatment (Fig. 4E). The increased stability of MYC-DR compared to MYC-WT in cells treated with cycloheximide (Fig. 4F) further indicates that site-specific deacetylation of MYC by HDAC2 in the CoREST complex stabilizes MYC by protecting it from proteolytic degradation in melanoma.

### Transcriptional control of DNA replication and mitosis by CoREST complex in melanoma cells

To gain insights into the functional roles of the CoREST complex in cancer cells, we performed transcriptomic analysis by RNA sequencing on three melanoma cell lines (A375, WM983B, and SK-MEL2) following RCOR1 knockdown (RCOR1-KD). Differentially expressed genes (DEGs) were identified by comparison with isogenic control cells. We found that decreased RCOR1 expression led to a greater number of upregulated genes than downregulated genes, consistent with the established function of CoREST complex as a transcriptional repressor.

However, a substantial number of genes were also downregulated in all three cell lines, suggesting that the CoREST complex may also play a role in the transcriptional activation of a subset of genes (Fig. 5A), which is aligned with other previous reports that LSD1 and MYC co-occupy proximal promoter region of a subset of genes and co-operatively transactivate transcription in embryonic stem cells (*33*).

**Fig. 5.**
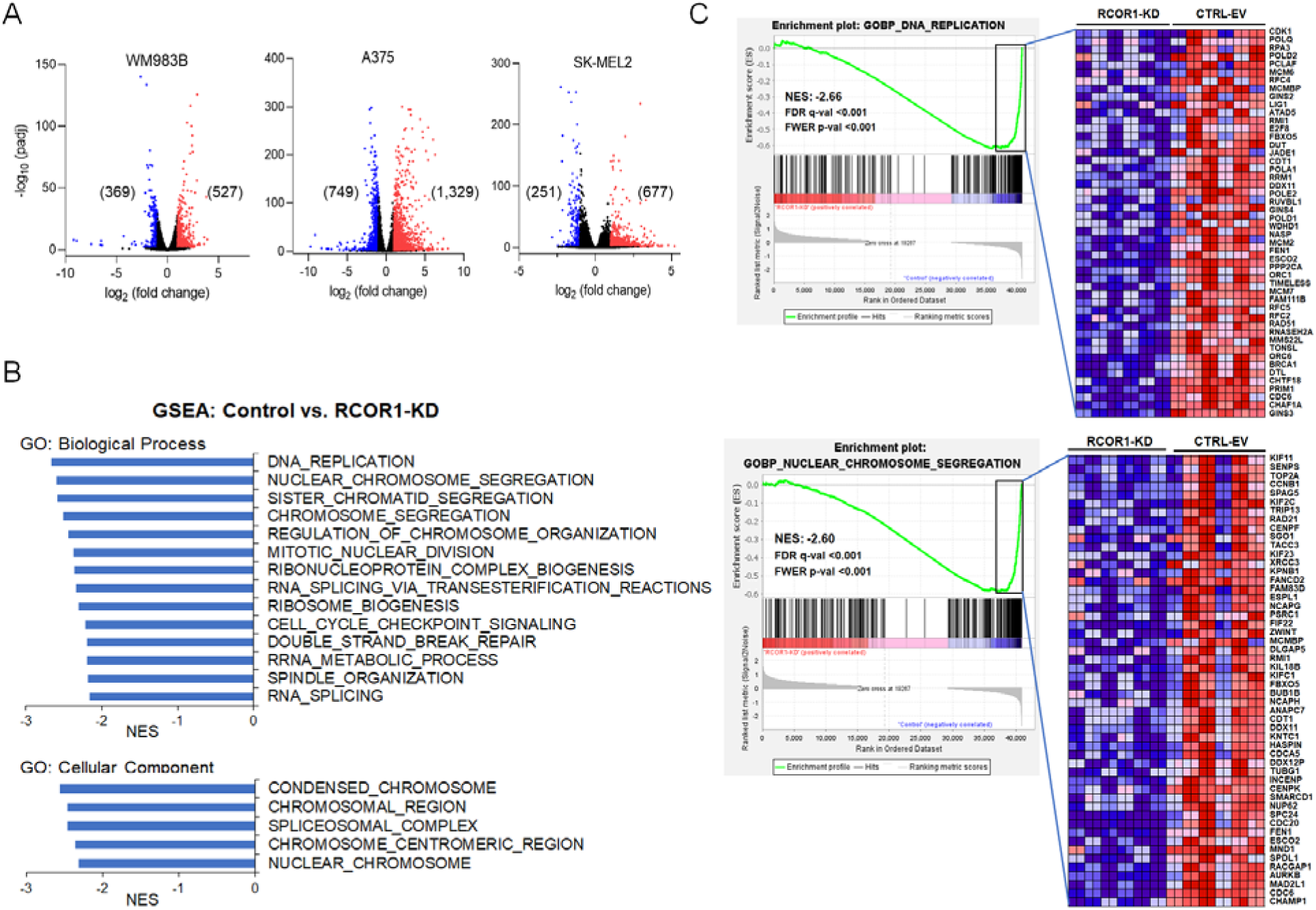
The CoREST complex transcriptionally upregulates genes functionally associated with DNA replication and mitotic chromosome segregation in melanoma. (A) Differentially expressed genes (DEGs) in melanoma cells with RCOR1-KD compared to isogenic control cells. (B) Gene Set Enrichment Analysis (GSEA) showing the top enriched categories for genes differentially expressed by RCOR1-KD in three melanoma cell lines. (C) GSEA plots and heatmaps of genes significantly downregulated in RCOR1-KD cells. The top two gene sets (GOBP: DNA Replication and GOBP: Nuclear Chromosome Segregation) are shown.

Gene ontology analysis using Gene Set Enrichment Analysis (GSEA) in conjunction with the human Molecular Signature Database (MSigDB v2023.1.Hs) was performed to identify gene signatures commonly regulated by the CoREST complex in these melanoma cell lines. While several gene sets associated with inflammation and immune responses were enriched among the upregulated genes (fig. S2), the most highly enriched gene sets among the downregulated genes fell into two functionally distinct groups. One group consisted of gene sets related to chromosome regulation, such as DNA replication and chromosome segregation. The other group comprised gene sets associated with RNA processing (Fig. 5B).

Significantly downregulated genes in the DNA replication gene set (Human Gene Set: GOBP_DNA_Replication) included many genes involved in DNA replication, particularly components of the replication fork origin firing complex, such as *ORC1*, *ORC6*, *MCMBP*, *MCM2*, *MCM6*, *GINS1*, *GINS2*, *GINS3*, and *CDC45* (Fig. 5C, upper panel, and table S1). Genes involved in the DNA double-strand break (DSB) repair process, especially those at sites of stalled or collapsed replication forks, such as *MMS22L*, *TONSL*, and *RAD51* (*34–36*), were also significantly downregulated in RCOR1-KD cells (Fig. 5C and table S1). These findings suggest that the CoREST complex plays a role in promoting DNA replication fork formation and firing by transcriptionally regulating key genes involved in that process.

RCOR1-KD in melanoma cells also led to significantly decreased expression of genes regulating mitotic chromosome segregation, including *MAD2L1*, *BUB1B*, *AURKB*, *CDC20*, *UBE2C* (*37–40*), and several kinesin family members (*KIFC1*, *KIF18B*, *KIF23*, *KIF2C*, and *KIF11*), motor proteins playing key spindle function (*41*) (Fig. 5C, lower panel, and table S2). These transcriptomic analyses indicate that one important roles of the CoREST complex in melanoma cells is to maintain chromosome stability by ensuring accurate chromosome segregation through spindle assembly, chromosome alignment, and microtubule dynamics. Furthermore, in patient tissue samples (ICGC/TCGA Pan-cancer and cutaneous melanoma), the expression levels of these genes are positively correlated with RCOR1 expression, suggesting that CoREST-mediated regulation of genome and mitotic chromosome stability is not limited to melanoma (fig. S3).

### Functional interplay between CoREST complex and MYC to maintain genome stability in cancer cells

To determine whether MYC mediates the CoREST complex-induced transcriptional activation of genes regulating DNA replication and chromosome segregation in melanoma cells, we performed RT-PCR analysis in melanoma cells following RCOR1 knockdown (RCOR1-KD), MYC knockdown (MYC-KD), and RCOR1-KD in the presence of simultaneous exogenous expression of a degradation-resistant MYC mutant (MYC-DR) (Fig. 6A). Candidate genes were selected from the gene sets GOBP: DNA Replication and GOBP: Nuclear Chromosome Segregation, representing two functional categories: replication factors involved in origin firing and mitotic factors involved in chromosome segregation (Fig. 5C). As expected, the expression levels of these genes were significantly decreased in both RCOR1-KD and MYC-KD cells, albeit less effectively in the latter. However, their expressions were restored by exogenous introduction of MYC-DR (Fig. 6B).

**Figure 6.**
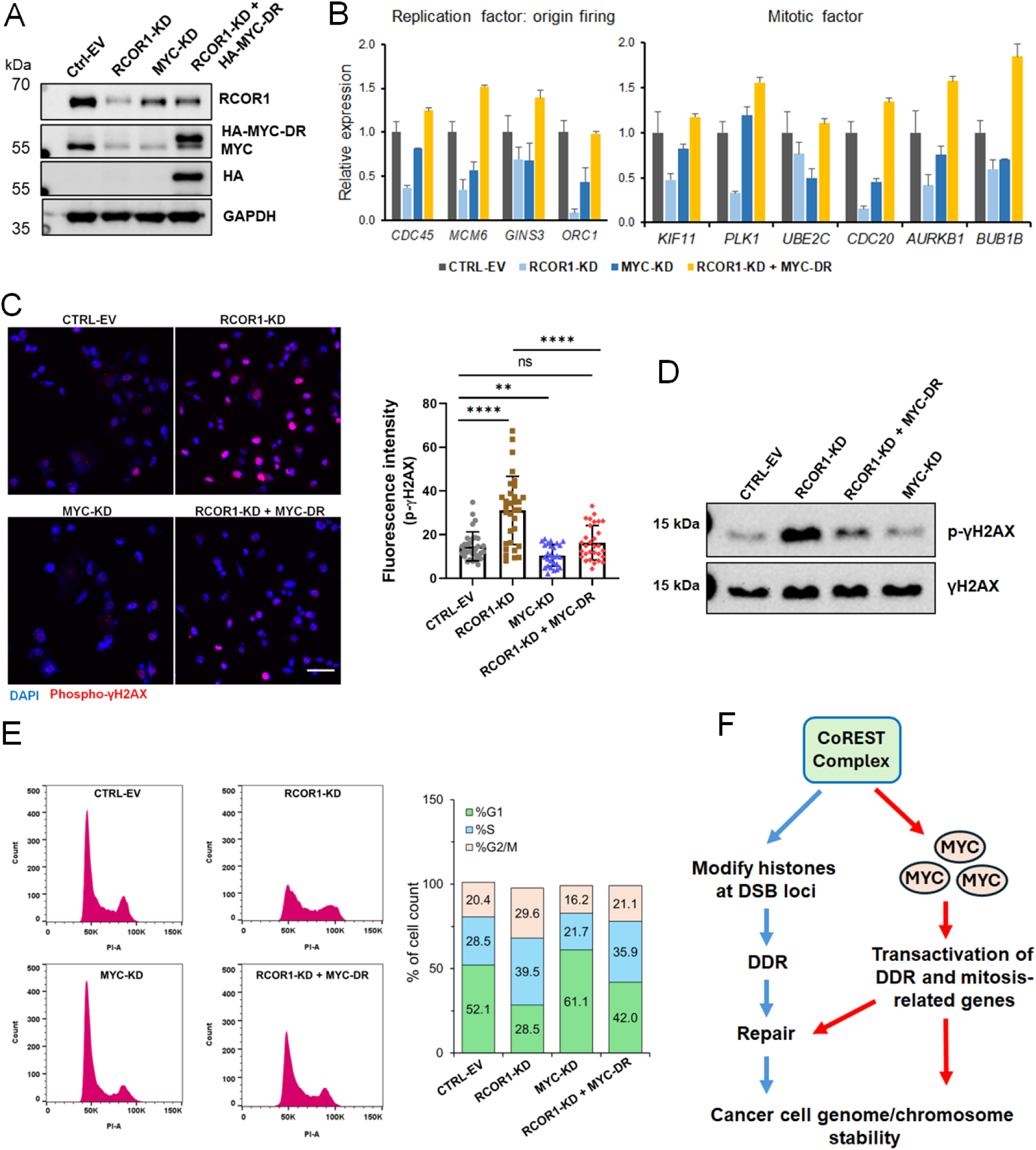
Oncogenic MYC mediates CoREST complex-induced chromosome stability in melanoma cells. **(A)** Immunoblots of parental A375 cells, RCOR1-KD cells, and RCOR1-KD cells with exogenous expression of HA-tagged MYC-DR. **(B)** RT-PCR analysis of genes selected from the gene sets for DNA replication and chromosome segregation (Figure 5C) in A375 cells with RCOR1-KD, MYC-KD, and RCOR1-KD + MYC-DR. **(C)** Immunofluorescence staining images of phospho-γH2A.X in A375 cells with RCOR1-KD, MYC-KD, and RCOR1-KD + MYC-DR. Scale bar: 50 μm. Student t-test, **, p<0.01, ****, p<0.0001, ns, non-significance. **(D)** Immunoblot assay of γH2A.X phosphorylation levels in A375 cells. **(E)** Cell cycle profiles by DNA content, as determined by flow cytometry, of A375 cells with RCOR1-KD and RCOR1-KD + MYC-DR. Cells were fixed with ethanol, and DNA content was measured using propidium iodide. **(F)** Schematic model illustrating the CoREST complex has dual roles in maintaining cancer cell genome stability by transcriptionally activating DDR-and mitosis-related genes and facilitating DDR by modulating histones at DSB loci in cancer cells.

The critical role of the CoREST complex in facilitating DNA replication and mitotic chromosome segregation by transcriptionally activating related genes in cancer cells was further validated by measuring DNA double-strand breaks (DSBs) and changes in DNA content during the cell cycle. A375 cells with RCOR1-KD exhibited markedly increased DSBs, as measured by phosphorylation of γH2A.X (*42*), while exogenous expression of MYC-DR reversed this phenotype. Cells with MYC depletion showed decrease DSBs compared to the control cells, indicating that the A375 cancer cells have an elevated basal level of replication stress (Fig. 6C). Immunoblot assays measuring the phosphorylation of γH2A.X also confirmed the effect of RCOR1 depletion (Fig. 6D). Cell cycle profile analysis further confirmed the genome-protective role of the CoREST complex, as an increased proportion of RCOR1-KD cells were arrested in the S and G2/M phases compared to control cells (Fig. 6E), a pattern typically observed in cells with DSB-induced CHK1/ATR checkpoint activation (*26, 43, 44*). These RCOR1-KD-induced phenotypes were rescued by exogenous expression of MYC-DR protein (Fig. 6E).

Pharmacological inhibition of the CoREST complex by Corin treatment of A375 cells also resulted in cell cycle arrest in the S and G2/M phases (fig. S4).

These experimental results strongly support the existence of a CoREST complex-MYC axis in cancer cells and indicate that one of the critical roles of the CoREST complex is to maintain and/or promote chromosome stability through upregulation of MYC expression. This mechanism enables cancer cells to sustain a high proliferative capacity while minimizing the risk of DNA damage-induced mitotic defects (Fig. 6F).

## Discussion

Our findings reveal a novel mechanism by which the CoREST complex, specifically through the HDAC1/2 subunit, stabilizes MYC protein by site-specific deacetylation of key lysine residues. This post-translational modification protects MYC from proteasomal degradation, thereby sustaining its oncogenic activity in melanoma cells. This result appears to contrast with earlier studies reporting that the acetylation of MYC can also enhance MYC stability and transcriptional activity (*22, 45*). For example, Hurd et al. (2023) showed that acetylation at K323 (a GCN5 target) increases MYC stability, while acetylation at K148(149) and K157(158) (p300 targets) does not confer similar stability and may even mark MYC for degradation(*45*). Our mutagenesis data, which identified five lysine residues critical for CoREST-mediated stabilization, suggest that deacetylation at these sites by HDAC2 is uniquely protective in melanoma cells.

Furthermore, this CoREST-mediated MYC stabilization mechanism may represent an adaptive response in cancer cells to maintain high MYC protein levels, supporting oncogenic proliferation and survival. Conversely, in other contexts or cancer types, acetylation might serve as a protective modification, depending on the specific lysines involved and the cellular milieu.

Taken together, this highlights the complexity of MYC regulation, where the functional outcome of acetylation is highly dependent on the specific residue, cellular context, and the repertoire of modifying enzymes present.

While the CoREST complex, comprising LSD1, HDACs, and RCOR proteins, is traditionally associated with transcriptional repression via histone demethylation (H3K4me1/2) and deacetylation, our findings reveal its unexpected role in transactivating a subset of MYC-dependent genes involved in DNA repair and mitotic regulation in cancer cells. This aligns with emerging studies demonstrating context-dependent transcriptional activation by CoREST complex. For instance, LSD1 and MYC co-occupy promoters of pluripotency genes in embryonic stem cells, where LSD1 facilitates MYC-driven transcription by resolving repressive chromatin loops, enabling RNA polymerase II recruitment (*33*). Similarly, in prostate cancer, LSD1 cooperates with MYC to activate metabolic genes such as *PHGDH*, with LSD1 depletion abolishing MYC’s transcriptional output (*46*). Our data extend these observations to melanoma, suggesting a conserved mechanism: CoREST complex may recruit the transcription factor MYC to the promoters of their target genes, which is supported by their direct interaction, thereby enabling sustained MYC activity.

Our data also suggest that the CoREST complex has a dual function in maintaining genome stability in cancer cells. First, the CoREST complex is recruited to sites of DNA double-strand breaks (DSBs), where it modulates local chromatin structure via histone deacetylation and demethylation, facilitating the recruitment of DNA repair machinery (*10, 11*). This is consistent with recent studies showing that CoREST interacts with DSB-associated factors such as USP22 and GSE1 to promote a chromatin environment conducive to efficient repair (*15*). Second, our transcriptomic analyses reveal that the CoREST complex, through stabilization of MYC, transcriptionally activates a suite of genes involved in DNA replication, DSB repair, and mitotic chromosome segregation. This dual role is particularly significant in cancer cells, which often experience heightened genome instability due to oncogenic MYC overexpression. By both initiating chromatin remodeling at DNA lesions and upregulating DDR and mitotic genes, the CoREST complex acts as a central coordinator of genome maintenance in the face of oncogenic stress. This may provide a survival advantage for cancer cells, enabling them to tolerate and repair increased DNA damage, thus promoting tumor progression.

Despite these advances, several limitations warrant consideration. While our data strongly support a direct role for HDAC2 in MYC deacetylation and stabilization, the precise molecular mechanism, such as the interplay between acetylation, ubiquitination, and degradation pathways, remains incompletely defined. Our study also leaves a critical mechanistic question unresolved. While RCOR1 knockdown (RCOR1-KD)-induced defects in DNA replication and mitotic progression were rescued by MYC-DR (degradation-resistant MYC), this does not fully distinguish whether the CoREST complex and MYC act independently, sequentially, or cooperatively to activate CoREST complex target genes involved in the DNA replication and mitotic chromosome segregation in cancer cells. The phenotype rescue assays suggest MYC is the dominant downstream effector for transcriptional activation of DNA repair and mitotic genes. However, we cannot exclude the possibility that the CoREST complex primes chromatin accessibility at these loci (e.g., via LSD1-mediated H3K4me2 demethylation or HDAC-mediated deacetylation) to facilitate MYC binding, implying a cooperative mechanism. For example, MYC may rely on CoREST-dependent chromatin remodeling to access promoters, even if MYC itself drives transcription. ChIP-seq experiments comparing CoREST and MYC occupancy in RCOR1-KD cells with/without MYC-DR could clarify this relationship.

## Methods

### Cell culture

The cell lines used in this study were cultured in DMEM supplemented with 10% FBS and 1% penicillin/streptomycin at 37°C with 5% CO2 in a humidified incubator. WM983B and 1205Lu cells were obtained from M. Herlyn (The Wistar Institute). All other cells were purchased from ATCC and tested to be free of mycoplasma.

### Plasmid construction for stable cell line generation

The knockdowns were performed using lentiviral plasmid pLKO.1-TRC. The plasmid was digested with EcoRI (NEB, USA) and AgeI (NEB, USA). The annealed shRNA oligos for the target genes were cloned into the EcoRI and AgeI sites of pLKO.1 vector and ligated with Ligase mixture (TAKARA, Japan). The shRNA clones were confirmed by Sanger sequencing.

For the exogenous expression of proteins, the genes were PCR amplified from cDNA, digested with BamHI (NEB, USA) and NotI (NEB, USA), and cloned into pCDH-EF1α-puro vector under the EF1α promoter and puromycin selection marker with HA or FLAG tag. The clones were confirmed by Sanger sequencing (GENEWIZ, NJ USA). The RCOR1 clone was purchased from a commercial company (U0745, GeneCopoeia).

The mutant MYC clones were amplified with primers containing the mutant amino acid sequences from the WT MYC clones and confirmed by Sanger sequencing (GENEWIZ, NJ USA). For the construction of lentiviral particles, Lenti-X-293T cells were transfected with ORF or shRNA vectors with packaging plasmids Pax2, PMD2.G at 3:2:1 using Lipofectamine 2000 (Invitrogen, USA). The viral particles were harvested 48 and 72 hrs post-transfection and filtered through a 0.45µm filter. Cells were transduced with viral particles and selected with 1µM puromycin for at least 72 hrs.

### RNA sequencing and analysis

Total RNA was extracted using TRIzol reagent from the melanoma cells (A375, WM983B, and SK-MEL2), stable RCOR1 knockdown and control, as well as Corin-treated and DMSO control cells. RNA was processed with the Ribo-Zero rRNA Removal Kit (Illumina) and further processed using TruSeq RNA Sample Prep Kit (Illumina). Sequencing libraries were prepared following the manufacturer’s recommendations. Samples were barcoded and sequenced using standard Illumina chemistry at GENEWIZ. The sequence reads were trimmed to remove possible adapter sequences and nucleotides with poor quality using Trimmomatic v.0.39. The trimmed reads were mapped to the Homo sapiens GRCh39 reference genome available on ENSEMBL using STAR aligner v.2.7.1b. Gene hit counts and unique gene hit counts were calculated using FeatureCounts from the Subread package v.2.0.3. Gene counts were further analyzed to test for differential expression using R package DESeq2 contrasting by RCOR1 KD or Corin treatment. For GSEA analysis, the differentially expressed genes (DEGs) between RCOR1 knockdown and control in each cell line were chosen up to 10 reads, and log2 fold changes between the samples were ranked from top to bottom. The dataset was analyzed using the software program GSEA4.3.2 downloaded from Broad Institute, and the enriched gene sets were identified based on human Molecular Signature Database (MSigDB v2023.1.Hs).

### Proximity ligation assay

PLA (DuoLink, Sigma) was performed according to the manufacturer’s instructions. Briefly, cells were fixed with 4% PFA for 10 min followed by permeabilization with 0.5% T-X-100 for 10 min. Cells were blocked in PLA blocking buffer for 30 min at 37°C and incubated with primary antibodies diluted in antibody dilution buffer for 1h at RT. Cells were washed twice in wash buffer A for 5 min and then incubated with PLA probes (rabbit (+) and mouse (-)) for 1h at 37°C. After washing twice with wash buffer A, the probes were ligated for 30 min at 37°C. The ligated product was amplified at 37°C for 100 min for fluorescent labeling. Cells were washed with wash buffer B for 10 min each followed by a brief wash with buffer B. Cells were mounted using In Situ Mounting Medium with DAPI, and the coverslip on the slide was sealed with nail polish. Images were acquired with the NIS-Elements Viewer microscope.

### Cell viability assay

Cell growth assay was performed using CellTiter-Glo 2.0 reagent (Promega, USA) according to the manufacturer’s instructions. Briefly, cells were grown in 96-well plates and a volume of CellTiter-Glo 2.0 reagent was added. The content was mixed on an orbital shaker for 2 min to lyse the cells and then incubated for 10 min at room temperature to stabilize the luminescent signal. The signal was acquired in 96 wells plate using a plate reader (TECAN, USA).

### Apoptosis assay

Cultured cells were trypsinized and washed two times with cold BioLegend Cell Staining Buffer and then resuspend with 500 ul Annexin V Binding Buffer. Add 5ul of FITC-conjugated Annexin V. Then, cells were stained with a viability dye, 7-AAD. After staining and washing, cells were acquired on a LSRFortessa flow cytometer (BD, USA) and analyzed using Diva software (BD, USA).

### Cell cycle profile assay

For cell cycles analysis, cells were trypsinized, washed with PBS and fixed in 70% ethanol on ice for 30 minutes. The ethanol was removed by centrifuging the cells for minutes at 6000 RPM. The pellet was washed with PBS two times. To the cell’s pellet RNAse A (100µg/ml) (Thermo Scientific, USA) were added followed by 400ul propidium iodide (PI) (50 µg/ml) solution and incubated for 10min at room temperature. The fluorescence intensity of the PI-DNA complex in the cell suspension was measured using a BD LSR Fortessa cell analyzer (405, 488, 561, 640nm lasers). For the analysis of cell cycle distribution, appropriate gating was initially established to eliminate cell debris (dead cells) and multi-cellular aggregates (cell clumps), followed by an examination of the distribution of individual cells across various stages of the cell cycle, as estimated based on the Watson-pragmatic model available in the FlowJo^Tm^ software package (Version 10, LLC USA).

### Recombinant protein expression in E. coli

For protein expression in E. coli BL21 DE3, a single colony was grown overnight at 4°C as a starter culture. The starter culture was transferred to a larger volume of LB media until the OD reached 0.4∼0.6 (∼3 hrs). For the induction of protein expression, 1mM IPTG was added to the media and incubated for at least 3 hrs. The cells were centrifuged at 4000 RPM for 10 min and the pellet was stored at-30°C until use. For the isolation of inclusion bodies (IBs), the pellet was resuspended in xFactor buffer (Takara, USA) and rotated at 4°C for 30 min, and the IBs were recovered by centrifugation at 12000 RPM for 20min. The pellet was again resuspended in lysis buffer (8M urea, 50mM sodium phosphate monobasic, 300mM sodium chloride (NaCl), 10mM imidazole, and 10% T-X-100, pH8) and sonicated 6 times for 15 sec ON and 15 sec OFF with 30 Amplitude. The sonicated cells were centrifuged for 30 min at 12000 RPM and the supernatant collected. The Ni-NTA beads were washed with the lysis buffer in a column and the cell lysate was added to the resin. The cell lysate was allowed to drain from the column by gravity. The resin was washed with a wash buffer (8M urea, 50mM sodium phosphate monobasic, 300mM NaCl, 20mM Imidazole, pH8).

The protein was eluted with an elution buffer (8M urea, 50mM sodium phosphate monobasic, 300mM NaCl, 250mM Imidazole, pH8). To remove the imidazole and refold the protein for downstream applications, the protein was dialyzed (Slide-A-Lyzer Dialysis Cassettes, Thermo Fisher, USA) against decreasing concentrations of urea and finally against 50mM Tris and 150mM NaCl buffer. The concentration of protein was measured with the BCA method and stored at-80°C in the storage buffer (50mM Tris, 150mM NaCl, 5mM DTT, 20% glycerol).

### Co-IP and immunoblotting

Cells were lysed with 1X non-denaturing lysis buffer (20mM Tris, 137mM NaCl, 1% NP-40, 0.2mM EDTA) and rotated for 30 min at 4°C. The cell lysate was centrifuged at 13,000 RPM for 15 min and supernatant was collected. The soluble fraction was incubated with indicated antibodies at 4°C overnight followed by incubation with A/G magnetic beads (Fisher Scientific, USA) for at least 4 hours. The immunoprecipitates were washed 5 times with lysis buffer and eluted in 2X SDS buffer by boiling for 10 min at 100°C. For western blot, the cells were lysed in RIPA buffer (10mM Tris, 1mM EDTA, 0.5mM EGTA, 1% T-X-100, 0.1% sodium deoxycholate, 0.1% SDS and 140mM NaCl) followed by incubation at 4°C with overhead rotation. The cell debris was removed by centrifugation. The histone proteins were extracted by resuspending the cell’s pellet in TEB buffer (PBS containing 0.5% Triton-X-100 (v/v), 2mM PMSF, 0.02% (w/v) NaN3) for 10 min on ice followed by centrifugation at 2000rpm for 10 min. The pellet was washed two times with TEB buffer and resuspended in 0.2N HCl overnight at 4°C. The samples were centrifuged at 2000 rpm for 10 min. The protein was quantified by BCA assay (Fisher Scientific, USA). Protein was resolved on SDS-PAGE and immunoblotting was carried out with indicated antibodies.

### RT-PCR

RNA was isolated by TRIzol reagent (Life Technologies, USA) according to the manufacturer’s instructions followed by DNase I (Invitrogen, USA) treatment. RNA was purified by phenol/chloroform and dissolved in nuclease-free water. RNA was reverse transcribed by SuperScript III first-strand synthesis system (ThermoFisher Scientific, USA) or cDNA synthesis Kit (Applied Biosystems, USA). The cDNA was subjected to semiquantitative RT-PCR using SYBR green master mix (TOYOBO, Japan). The sequences of primers used in the PCR are listed in table S3.

### Immunofluorescence

Cells were fixed in 4% PFA (MP Biomedicals, USA) in PBS for 10 min and permeabilized with 0.5% T-X-100 for 10 min at room temperature. The permeabilized cells were incubated with blocking buffer (1% BSA in 0.1% T-X-100) for 1 hour followed by incubation with primary antibody overnight. Cells were washed 3 times with 0.5% T-X-100 and incubated with appropriate Alexa Fluor secondary antibodies (Invitrogen, USA). Images were captured with an inverted fluorescence microscope (Nikon).

### Statistics and reproducibility

Statistical analyses for group comparisons were performed using Student’s two-sided *t*-test and ANOVA, with significance defined as *p* < 0.05. All results are presented as the mean ± standard deviation (SD) from at least three biological replicates.

Analyses were conducted using GraphPad Prism software (version 9.5.1).

### Chemicals

Corin was synthesized in the laboratory of P.A. Cole (Harvard Medical School). MS275, GSK2879552, propidium iodide, and P300/CBP inhibitor A-485 were purchased from MedChem Express (Monmouth Junction, NJ USA).

## Acknowledgments

The authors would like to express our sincere gratitude to Philip A Cole at Harvard Medical School and Brigham and Women’s Hospital for his invaluable contributions and research materials that greatly enriched this study.

## Author contributions

Conceptualization: AAK, BR. Methodology: AAK, SK, TS, JB, JS, BR. Investigation: AAK, SK, TS, JB. Visualization: AAK, SK, JB, JS, BR. Data curation: AAK, AAA, BR.

Funding acquisition: JS, BR. Supervision: JS, BR. Writing – original draft: BR.

Writing – review and editing: AAK, SK, TS, JB, AAA, PDD, JLZ, JS, BR

## Data and materials availability

Data availability: Raw RNA-seq files have been deposited in the National Center for Biotechnology Information BioProject (access code: PRJNA996569). Other raw data are available upon request from the corresponding author.

## Supplementary Materials

The PDF file includes:

Fig. S1 to S4

Legends for Tables S1 to S3

Other Supplementary Materials for this manuscript includes the following: Table S1 to S3

