## Supplemental data for "CoREST Complex Stabilizes MYC Protein to Promote Cancer Cell Genome Stability"

**Supplementary Data**

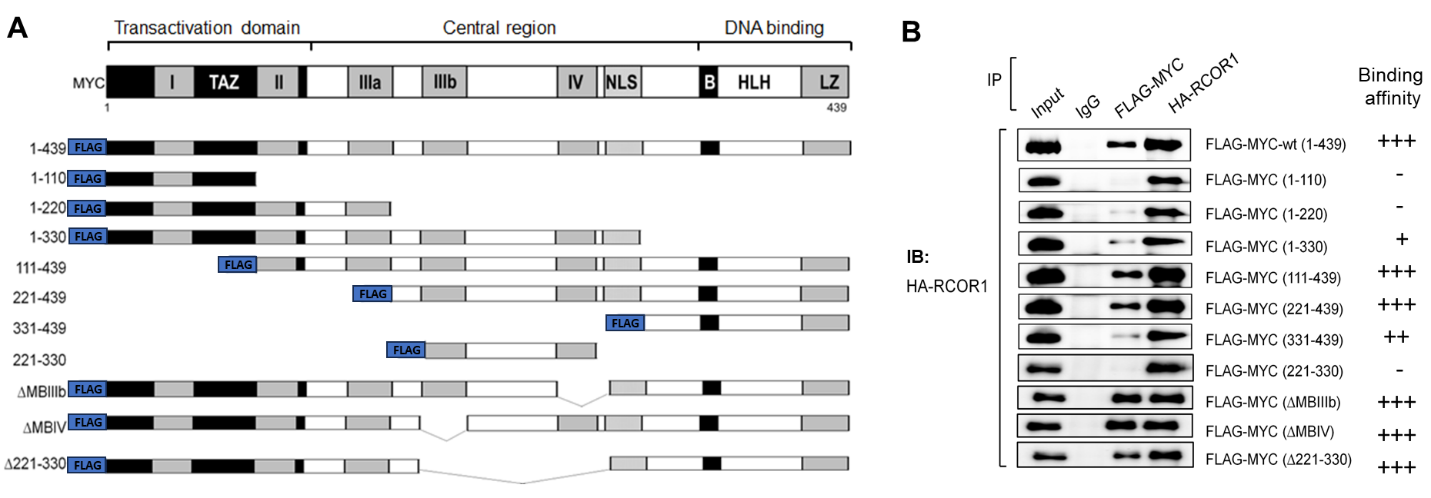

**Figure S1. Interaction domain mapping in MYC for CoREST-complex. A)** Protein interaction domains of MYC and 3xFLAG-tagged deletion mutants. **B)** Co-immunoprecipitations of deletion mutants of MYC with RCOR1 in HEK293T cells. 3xFLAG-tagged MYC deletion mutants and HA-tagged full-length RCOR1 expressed in HEK293T cells were pulled down and blotted with anti-HA antibodies.

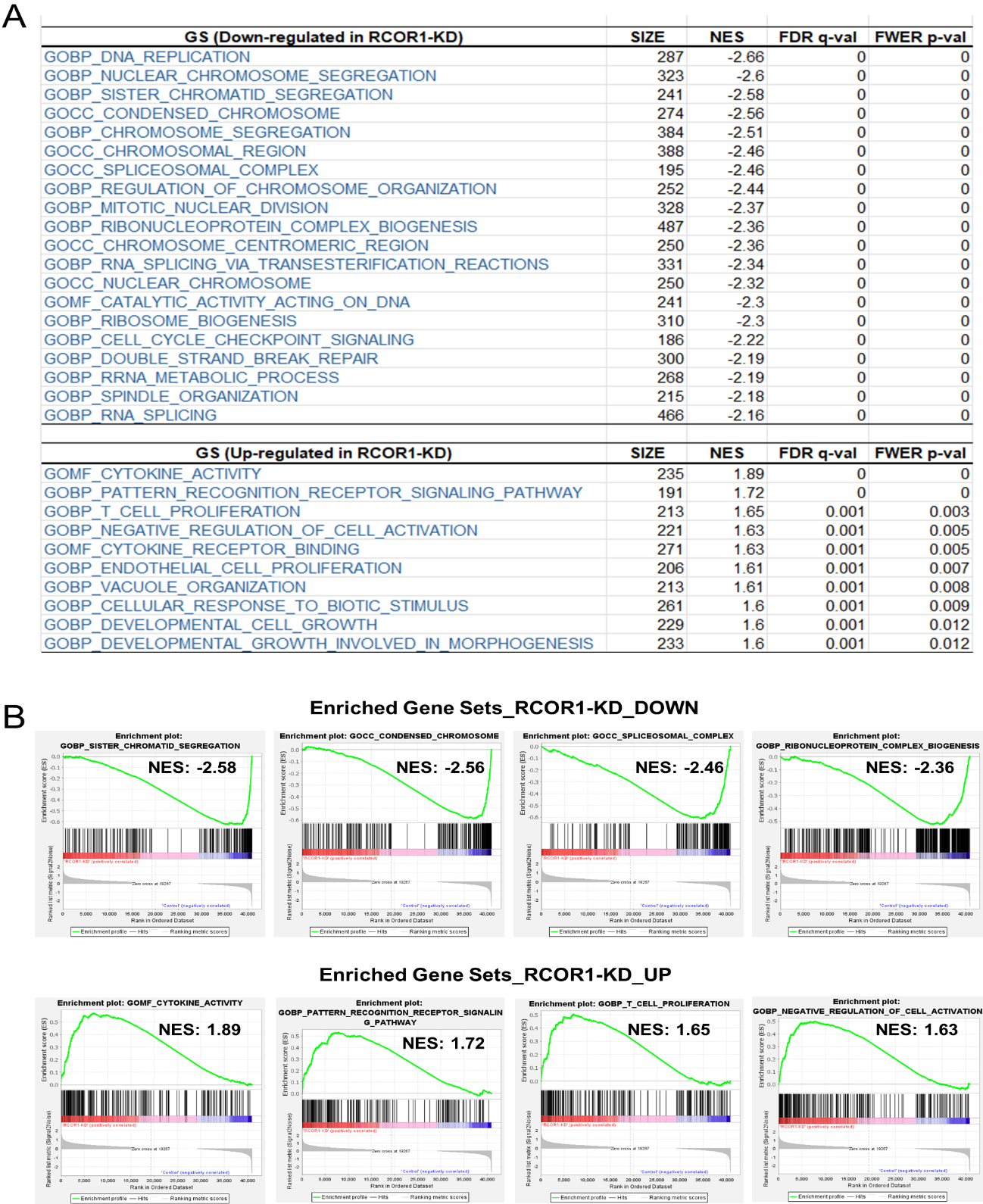

**Figure S2. Gene set enrichment analyses (GSEA).** Down-regulated (DOWN) and Up-regulated (UP) genes in three melanoma cell lines (WM983B, A375, and SK-MEL2) with RCOR1-KD vs control cells. **A)** Top deregulated gene sets. **B)** Selected enrichment plots with normalized enrichment scores (NES).

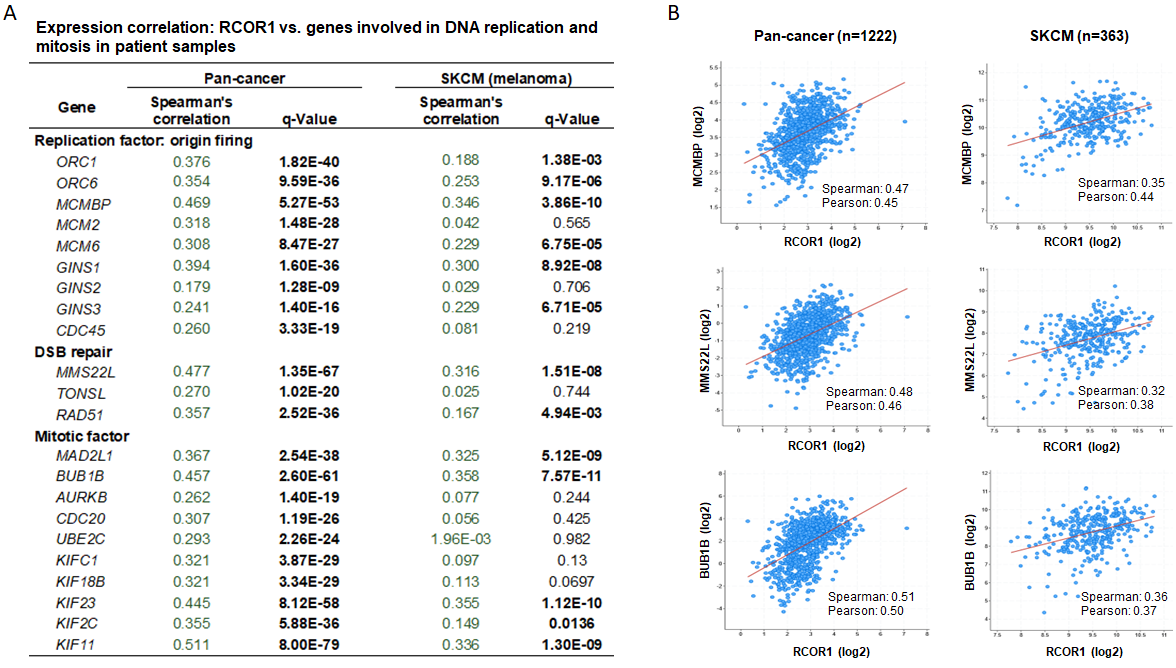
**Figure S3**.

**Figure S3. Expression correlations between RCOR1 and genes involved in DNA replication, DSB repair, and mitotic chromosome segregation in cancer patient samples. A)** Spearman’s correlations between expression of RCOR1 and significantly down-regulated genes that are selected from the gene sets of GOBP: RNA Replication and GOBP: Nuclear Chromosome Segregation in patient samples. For the analysis of pan-cancer patient samples, RNA-sequence data set of 1222 patient samples from ICGC (International Cancer Genome Consortium) were used. For the analysis of cutaneous melanoma (SKCM) samples, RNA-sequence data set of 363 samples from TCGA (The Cancer Genome Atlas) were used. **B)** Correlation plots of RCOR1 vs MCMBP, RCOR1 vs. MMS22L, and RCOR1 vs. BUB1B in pan-cancer and melanoma cohorts.

**Figure S4**

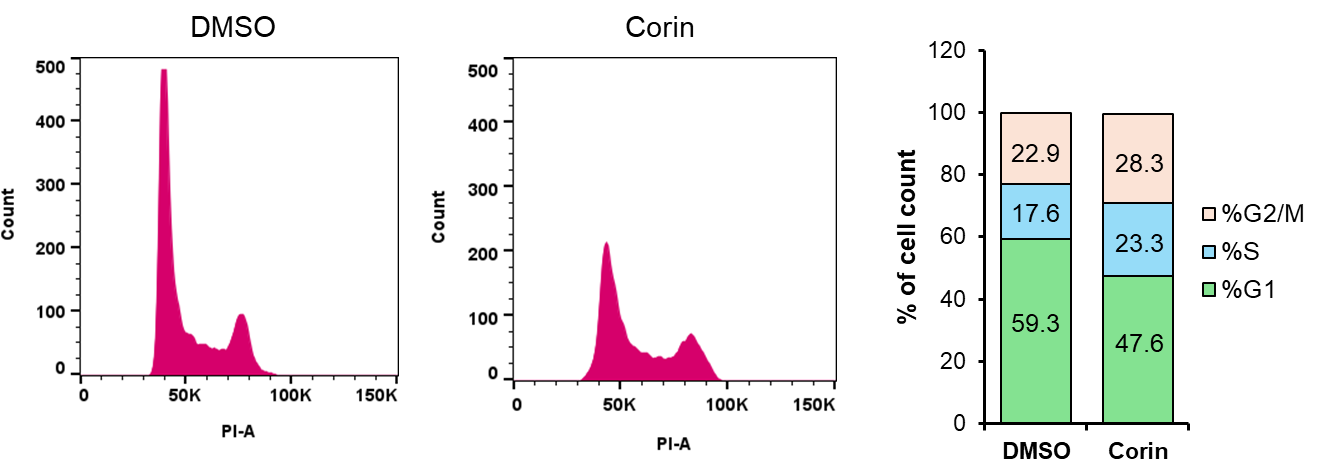

**Figure S4. Flow cytometry analysis of DNA content profiles in A375 cells treated with Corin. Cells were treated with Corin (2** μM) and DMSO for 48 hours. Following treatment, cells were fixed with ethanol and DNA content was measured using propidium iodide.

**Table S1. A gene list of Gene set: GOBP_DNA_Replicatiopn from the down-regulated genes in RCOR1-KD cells compared to control.** Genes with Running ES (enrichment score) equal or less than -0.6176 were identified as enriched genes. Yes: enriched, No: not enriched.

**Table S2. A gene list of Gene set: GOBP_Nuclear_Chromosome_Segregation from the down-regulated genes in RCOR1-KD cells compared to control.** Genes with Running ES (enrichment score) equal or less than -0.5845 are identified as enriched genes. Yes: enriched, No: not enriched.

**Table S3. Lists of antibodies and nucleotide sequences for RT-PCR assays and shRNA gene knockdown experiment used in this study**
